# Co-evolutionary dynamics of defense systems and anti-defense genes in chromosomes and mobile genetic elements of *Enterobacteriaceae*

**DOI:** 10.64898/2025.12.17.694813

**Authors:** Yang Liu, Stephen Dela Ahator, João Botelho, Jaime Iranzo

**Affiliations:** Department of Microbiology, Zhejiang University School of Medicine, Zhejiang, China; Drug Transport and Delivery Group, Department of Pharmacy, UiT─ The Arctic University of Norway, Tromsø, Norway; Universidade Católica Portuguesa, CBQF - Centro de Biotecnologia e Química Fina – Laboratório Associado, Escola Superior de Biotecnologia, Rua Diogo Botelho 1327, 4169-005 Porto, Portugal; Centro de Astrobiología (CAB, CSIC-INTA), Torrejón de Ardoz, Madrid, Spain

**Keywords:** bacterial immunity, anti-defense systems, gene gain and loss, mobile genetic elements, Enterobacteriaceae

## Abstract

Bacterial defense systems undergo rapid evolutionary changes, yet the specific contributions of the genomic compartment (chromosomal versus mobile genetic element [MGE]) to these dynamics remain unclear. We analyzed 6,907 genomes from 213 species of *Enterobacteriaceae* using a phylogenetic birth–death model to quantify gain, loss, expansion, and reduction of defense (DEF) and anti-defense (anti-DEF) gene families. We found that DEF turnover differs from the background dynamics of the genomic elements that carry them. Direct comparisons among DEFs further revealed differences in loss and copy-number dynamics across genomic compartments, with the highest rates on chromosomes, intermediate rates on plasmids and ICEs/IMEs, and the lowest rates on prophages, whereas the relative balance between gain and loss and between expansion and reduction remained similar across chromosomes and MGEs. MGE-associated anti-DEFs showed higher loss, expansion, and reduction rates than DEFs, whereas chromosome-associated anti-DEFs showed lower gain rates than DEFs. Finally, we found that gains and losses of DEFs and anti-DEFs were positively correlated across bacterial lineages, with positive associations observed for both matching (that is, antagonistic) and non-matching pairs. Together, these results show that genomic background affects the overall flux of DEFs and anti-DEFs, and that turnover of DEFs and anti-DEFs is evolutionarily coupled across lineages.

## INTRODUCTION

Mobile genetic elements (MGEs), such as bacteriophages and plasmids, play a dual role in evolution as vectors of horizontal gene transfer (HGT) and potentially damaging genomic parasites ^1–3^. To survive the constant pressure of MGEs, bacteria have evolved a vast and diverse arsenal of defense systems (DEFs), ranging from restriction-modification (RM) and CRISPR-Cas to recently discovered mechanisms like CBASS, Gabija, and various toxin-antitoxin modules ^4–8^. The distribution of these systems across bacterial genomes is highly uneven, consistent with rapid evolutionary turnover ^5,9^. Intriguingly, DEFs are frequently encoded within MGEs, which are the very elements they are thought to control ^4,10–15^.

Recent large-scale phylogenomic analyses have highlighted that the evolutionary impact of DEFs is shaped by their physical linkage with MGEs ^16–19^. For instance, we recently demonstrated that the inhibitory effect of defense systems on HGT is often masked by their co-transfer with MGEs, resulting in a positive association between defense presence and MGE abundance over short evolutionary timescales ^18^. This linkage implies that the evolutionary dynamics of DEFs may be linked to the life cycles of their genomic carriers. Yet, the precise nature of this dependency remains to be fully disentangled. While chromosomes generally provide a stable evolutionary background, plasmids and prophages are governed by high mobility and instability ^3,20,21^. It remains unclear whether DEFs residing on chromosomes and MGEs exhibit uniform evolutionary rates, or if they follow location-specific dynamics of gene gain, loss, expansion, and reduction.

Furthermore, the evolution of DEFs is driven by a continuous arms race with MGE-encoded anti-defense systems (anti-DEFs), such as anti-CRISPR (Acr) and anti-RM proteins ^22–26^. Anti-DEFs can suppress host immunity, facilitate MGE invasion and potentially alter the selection pressure on defense *loci*. While previous studies have established the prevalence of these counter-defense mechanisms ^23,27^, quantitative comparisons of their evolutionary turnover rates relative to their target DEFs are lacking. Specifically, it is unknown whether anti-defense genes essentially mirror the turnover of defense genes or if they exhibit accelerated dynamics indicative of a distinct evolutionary strategy.

In this study, we dissect the large-scale evolutionary dynamics of DEFs and anti-DEFs within *Enterobacteriaceae*, a clinically relevant family known for its extensive mobilome and diverse immune repertoire. By analyzing 6,907 high-quality genomes across 213 species using a phylogenetic birth–death model, we quantified the rates of gene gain, loss, expansion, and reduction for DEFs located on chromosomes and MGEs. We aimed to determine: (i) how the genomic compartment (chromosomal vs. MGE) contributes to the turnover bias of defense genes; (ii) whether DEFs and anti-DEFs exhibit coupled evolutionary trajectories; and (iii) how these dynamics correlate with broader functional gene categories.

## RESULTS

### DEF turnover differs from non-defense gene backgrounds

To evaluate whether the evolutionary dynamics of DEF genes differ from the dynamics of other genes from chromosomal and MGE backgrounds, we quantified the rates of gene family gain, loss, expansion, and reduction across 6,907 genomes from 213 species of *Enterobacteriaceae* (Supplementary Table S1) using a phylogenetic birth–death model implemented in the software Count. These four processes represent, respectively, the acquisition of new gene families, loss of entire families, increase in copy number within existing families, and copy-number reduction in multi-copy gene families (Figure 1A-B). As a preliminary step, we built orthologous gene families (OGs) for each species, identified those belonging to defense and anti-defense systems, and classified them as chromosome-associated, prophage-associated, satellite-associated, plasmid-associated, or integrative and conjugative/mobilizable element (ICE/IME)-associated based on their genomic location (the chromosome category excluded prophages and other known integrative MGEs, such as satellites, ICEs, and IMEs; see Methods and Supplementary Figure S1). Overall, we identified 14,506 chromosome-associated DEF gene families, representing the largest fraction of defense genes (Figure 2A). Among MGEs, prophages contained 897 gene families, followed by plasmids (698) and ICEs/IMEs (384). A substantial fraction of DEF gene families appeared both in the chromosome and one or more classes of MGEs (6,788). Sharing of DEF gene families across MGE classes was most often observed for plasmids and ICEs/IMEs (192 OGs) and, less frequently, for prophages and plasmids (80 OGs). Regarding anti-DEF gene families, the highest number was found in plasmids (597), followed by chromosomes (320) and prophages (212), with overlap observed between chromosomes and plasmids (341 OGs) and between chromosomes and prophages (243 OGs) (Figure 2B).

**Figure 1.**
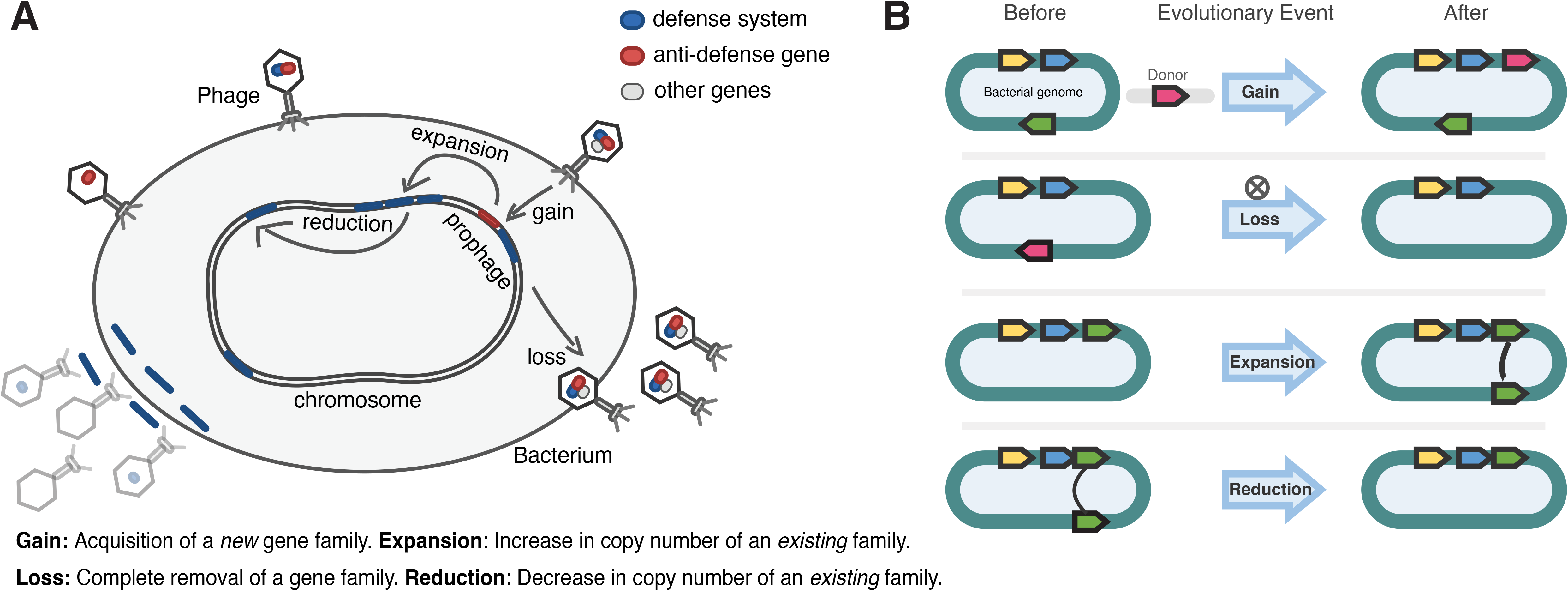
Schematic overview of evolutionary turnover processes affecting defense systems and anti-defense genes. (A) Conceptual illustration of defense systems (DEFs), anti-defense genes (anti-DEFs), and other genes within a bacterial genome, showing gene turnover associated with the chromosome and prophage regions. (B) Schematic definitions of the four evolutionary processes analyzed: gain, acquisition of a new gene family; loss, complete removal of a gene family; expansion, increase in copy number of an existing gene family; and reduction, decrease in copy number of an existing gene family.

**Figure 2.**
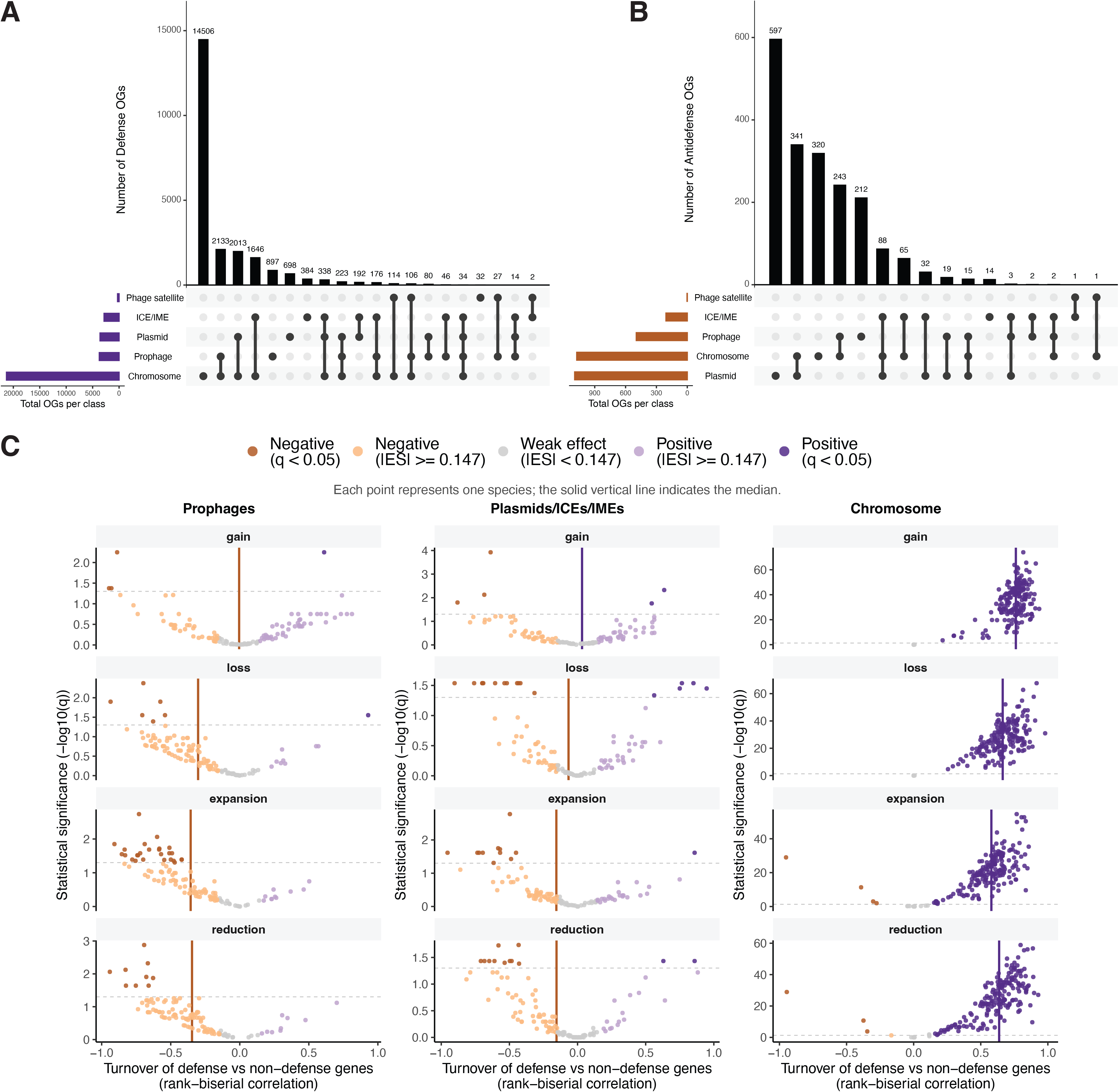
Genomic distribution of DEF and anti-DEF orthologous groups and their turnover relative to local non-defense genes. (A–B) Distribution of DEF (A) and anti-DEF (B) orthologous groups (OGs) across chromosomes, prophages, plasmids, ICEs/IMEs, and phage satellites. OGs were assigned using a 90% classification threshold. Connected dots indicate the genomic-element classes contributing to each combined category. (C) Differences in evolutionary rates between DEFs and non-defense genes (ND) located within the same genomic background, shown for prophages (pDEF vs ND), plasmids plus ICEs/IMEs (pl+iDEF vs ND), and chromosomes (cDEF vs ND). Each point represents one *Enterobacteriaceae* species. The x-axis represents the rank-biserial correlation effect size (ES); the y-axis represents statistical significance as −log10(q). Colors indicate statistical significance and effect-size categories using |ES| ≥ 0.147 as the primary effect-size threshold^63,64^. Solid vertical lines indicate the median ES across species, and the horizontal dashed line indicates q = 0.05. ES, effect size.

For all subsequent analyses, we focused on gene families associated with a single genomic compartment (the chromosome or a single class of MGE), as they better represent compartment-specific dynamics. We then used the presence-absence profiles of these families to infer the turnover rates of chromosomal DEFs (cDEF), prophage DEFs (pDEF), phage satellite DEFs (satDEF), and plasmid plus ICE/IME DEFs (pl+iDEF) and compared those to the turnover rates of non-defense (ND) genes located in the same genomic compartments (Methods). Comparisons between DEF and ND genes revealed distinct patterns across genomic compartments. In prophage regions, pDEF generally display lower loss, expansion, and reduction rates than the general (ND) prophage gene pool (Figure 2C, left; positive/negative effect size, defined as rank-biserial correlation beyond +/-0.147, in 45/40 (*P* = 0.665, n=120), 9/85 (n=120), and 10/94 (*P* < 0.001, n=119), and 9/93 (*P* < 0.001, n=119) species for gain, loss, expansion and reduction, respectively; binomial sign tests). These results suggest that pDEFs experience more stable retention and fewer copy-number fluctuations than other prophage genes. Qualitatively similar effect-size patterns, though weaker in magnitude, were observed for plasmids and ICEs/IMEs (Figure 2C, center; the number of species in which the effect size is positive/negative with a minimum rank-biserial correlation of +/-0.147, is 41/41 (*P* = 1.0), 33/48 (*P* = 0.119), 24/64 (*P* < 0.001), and 17/61 (*P* < 0.001) for gain, loss, expansion, and reduction, respectively; n = 118 for all comparisons, binomial sign tests). In stark contrast, on chromosomes, cDEFs showed substantially higher turnover rates than chromosomal ND gene families across all four event types, with extensive support from both *q* values and effect sizes (Figure 2C right; positive/negative effect sizes in 190/0 (n=196), 190/0 (n=195), 179/4 (n=195), and 182/4 (n=196) species for gain, loss, expansion, and reduction, respectively; all *P* < 0.001, binomial sign tests). Phage satellite-associated DEFs were not included in subsequent statistical comparisons because too few species met the minimum sample-size requirements. We assessed the robustness of these results using alternative effect size thresholds of +/-0.33 and +/-0.474 across all meta-analyses, and the main conclusions remained consistent (Supplementary Figure S3). To more rigorously assess whether the patterns observed in the species-level comparisons were statistically supported, we fitted generalized linear mixed-effects models (GLMMs) with the functional class (DEF or ND) as predictor and species as random effects (Methods). In prophage regions, the DEF/ND rate ratios were 0.28, 0.01, and 0.03 for loss, expansion, and reduction, respectively, indicating lower rates for pDEFs than for pND genes (all *q* < 0.001; Supplementary Figure S4). The corresponding ratios in plasmid plus ICE/IME regions were 0.71, 0.08, and 0.26 (all *q* < 0.001; Supplementary Figure S4). In contrast, chromosomal DEFs showed higher gain, loss, and reduction rates than chromosomal ND genes (cDEF/cND rate ratios = 4.41, 2.78, and 1.34, respectively; all *q* < 0.005), whereas expansion did not differ significantly.

### DEFs show higher retention in MGE-associated genomic compartments

We next compared turnover dynamics directly among DEFs located on chromosomes, prophages, and plasmids plus ICEs/IMEs. GLMMs revealed significant differences in DEF turnover rates across genomic compartments (Figure 3A). cDEFs showed higher rates of loss, expansion, and reduction than pDEFs and pl+iDEFs (cDEF vs pDEF: 2.72x, 135.19x, and 24.59x for loss, expansion, and reduction, respectively; cDEF vs pl+iDEF: 1.50x, 33.04x, and 9.51x, respectively; all *q* < 0.001). pl+iDEFs showed higher rates than pDEFs (1.81x, 4.09x, and 2.59x for loss, expansion, and reduction, respectively; all *q* < 0.001). Overall, DEFs showed the highest turnover on chromosomes, intermediate turnover on plasmids plus ICEs/IMEs, and the lowest turnover on prophages. Gain rates showed weaker differences, with only the cDEF versus pl+iDEF comparison reaching statistical significance (1.13x; 95% confidence interval [CI], 1.03–1.24; *q* = 0.032). Moreover, the gain/loss and expansion/reduction ratio did not differ significantly among chromosomal and MGE-associated DEFs (Figure 3B). The MGE-to-chromosome ratio was 1.08 for gain/loss (95% CI, 0.98–1.20; *q* = 0.137), and 0.90 for expansion/reduction (95% CI, 0.80–1.01; *q* = 0.130).

**Figure 3.**
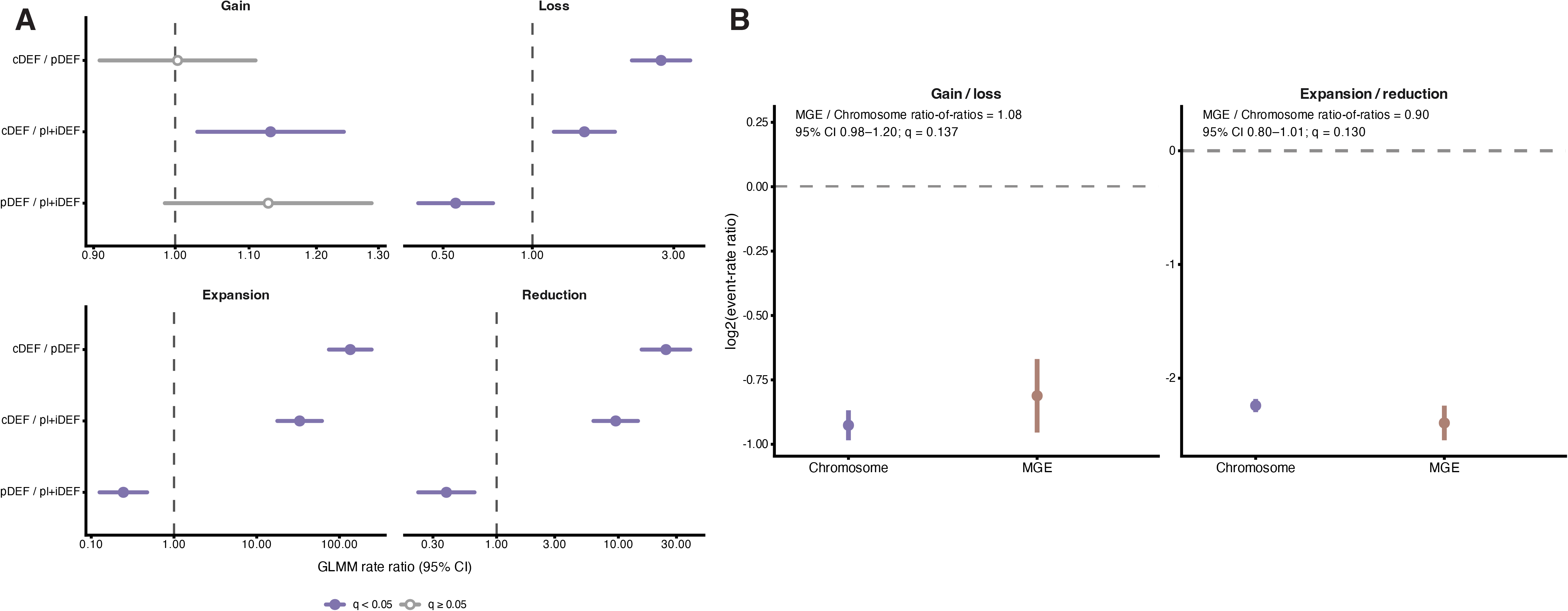
GLMM-based comparison of DEF turnover across genomic compartments. (A) Pairwise comparisons of DEF evolutionary rates among chromosomes (cDEF), prophages (pDEF), and plasmids plus ICEs/IMEs (pl+iDEF) for gain, loss, expansion, and reduction events. Points represent estimated rate ratios from generalized linear mixed models (GLMMs), and horizontal lines indicate 95% confidence intervals (CIs). The dashed vertical line indicates a rate ratio of 1. Filled points indicate q < 0.05 and open points indicate q ≥ 0.05. (B) Gain-to-loss and expansion-to-reduction log2-transformed rate ratios for chromosomal and MGE-associated DEFs. Points and error bars represent estimated ratios and 95% CIs.

### Anti-DEFs exhibit higher rates of loss and copy-number change than DEFs

We next used GLMMs to compare DEFs and anti-DEFs within the same genomic elements, including prophages, plasmids plus ICEs/IMEs, as well as chromosomes (Figure 4). DEFs showed significantly lower loss, expansion, and reduction rates than anti-DEFs in prophages and plasmids plus ICEs/IMEs (anti-DEFs showed 2.36x, 13.63x, and 5.51x higher loss, expansion, and reduction rates than DEFs in prophages, respectively, and 1.59x, 11.53x, and 5.59x higher rates in plasmids plus ICEs/IMEs; all *q* ≤ 0.003). No significant differences were detected between gain rates of DEFs and anti-DEFs in MGEs. In contrast, chromosomal DEFs exhibited higher gain rates than chromosomal anti-DEFs (11.41x; 95% CI, 5.07–25.64; *q* < 0.001), but similar rates of loss, expansion, and reduction. These results suggest that the higher loss and copy-number plasticity of anti-DEFs relative to DEFs is primarily associated with MGE genomic backgrounds.

**Figure 4.**
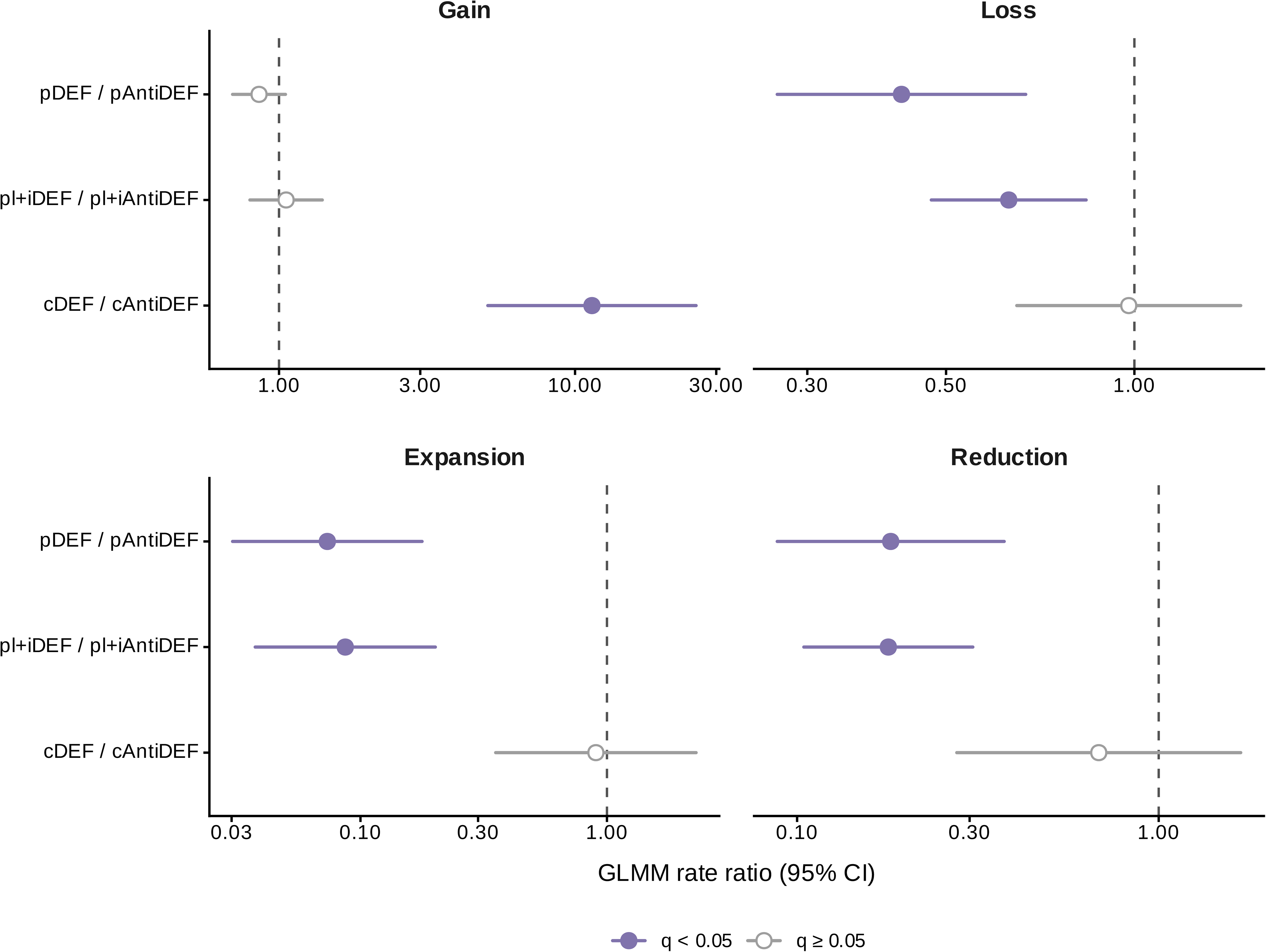
GLMM-based comparison of evolutionary turnover between DEFs and anti-DEFs across genomic compartments. Comparisons of DEF and anti-DEF evolutionary rates within prophages (pDEF/pAntiDEF), plasmids plus ICEs/IMEs (pl+iDEF/pl+iAntiDEF), and chromosomes (cDEF/cAntiDEF) for gain, loss, expansion, and reduction events. Points represent estimated rate ratios from generalized linear mixed models (GLMMs), and horizontal lines indicate 95% confidence intervals (CIs). The dashed vertical line indicates a rate ratio of 1. Filled points indicate q < 0.05 and open points indicate q ≥ 0.05.

To assess whether these trends are general or restricted to some specific defense systems, we performed a stratified analysis of copy number dynamics within the most widespread DEF and anti-DEF classes (Figure 5 and Supplementary Figure S5). For gain events, no statistically significant differences were detected between any DEF and anti-DEF pairs (Figure 5 and Supplementary Figure S5). In contrast, anti-CBASS and anti-RM systems showed higher loss, expansion, and reduction rates than their respective DEFs (anti-CBASS: *q* = 2.87 × 10^-5^, 0.0013, and 6.70 × 10^-5^; anti-RM: *q* = 0.0060, 4.48 × 10^-12^, and 1.03 × 10^-8^ for loss, expansion, and reduction, respectively; Wilcoxon signed-rank test; Figure 5 and Supplementary Figure S5). Four other classes (Retron/anti-Retron, Dnd/anti-Dnd, RloC/anti-RecBCD, and Thoeris/anti-Thoeris) show higher rates of anti-DEF loss and copy number change than their respective DEFs, although most comparisons were not statistically significant. An exception was the reduction rate of RloC/anti-RecBCD (*q* = 0.045). Taken together, these analyses confirm that the trends detected by the overall GLMM analysis (Figure 4) extend over multiple classes of DEFs and anti-DEFs, with significant differences detected in some of the most abundant systems.

**Figure 5.**
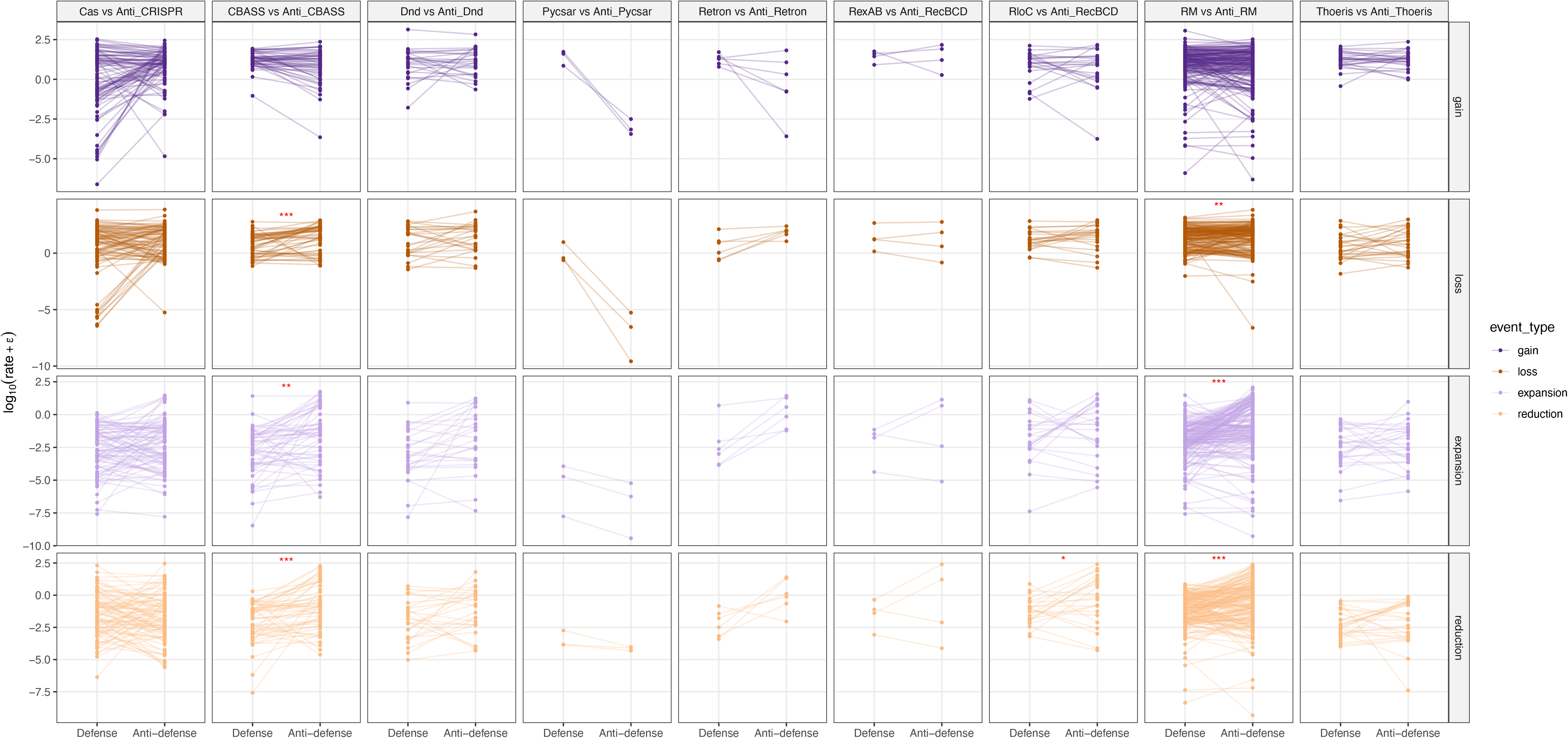
Paired evolutionary turnover rates of different classes of DEFs and anti-DEFs. Evolutionary rates of DEFs and anti-DEFs within the same species. Each line connects the median rate of a DEF with that of its corresponding anti-DEF in a single species. The y-axis represents log-transformed rates, where ε = 1 x 10^-16^. Red asterisks indicate statistically significant differences based on paired Wilcoxon signed-rank tests after multiple-testing correction (* q < 0.05, ** q < 0.01, *** q < 0.001).

### DEFs and anti-DEFs show coupled evolutionary turnover

To understand the evolutionary associations between DEFs, anti-DEFs, and broader functional gene categories, we calculated partial Spearman correlations for the number of evolutionary events per branch across all phylogenetic trees, controlling for branch length and using phylogenetic Brownian Motion simulations to assess statistical significance (Methods). Correlations were computed at the whole-genome level (including chromosomal plus MGE genes) and also separately for prophage-associated genes and plasmid-associated genes. We first examined general associations between DEF and anti-DEF turnover. For gene gain events, 23 of 191 species in the whole-genome analysis, 22 of 105 species in the prophage-specific analysis, and 48 of 112 species in the plasmid-specific analysis exhibited strong positive correlations with high statistical significance (r ≥ 0.5, *q < 0.05*; Figure 6A-C). Significant correlations were also observed for loss events (whole-genome: 117 out of 191 species; prophage-specific: 40 out of 105 species; plasmid-specific: 64 out of 112 species). Taken together, these analyses suggest that concerted gain and loss of DEF and anti-DEF occur at the whole-genome level and involve both prophage-associated and plasmid-associated DEFs and anti-DEFs.

**Figure 6.**
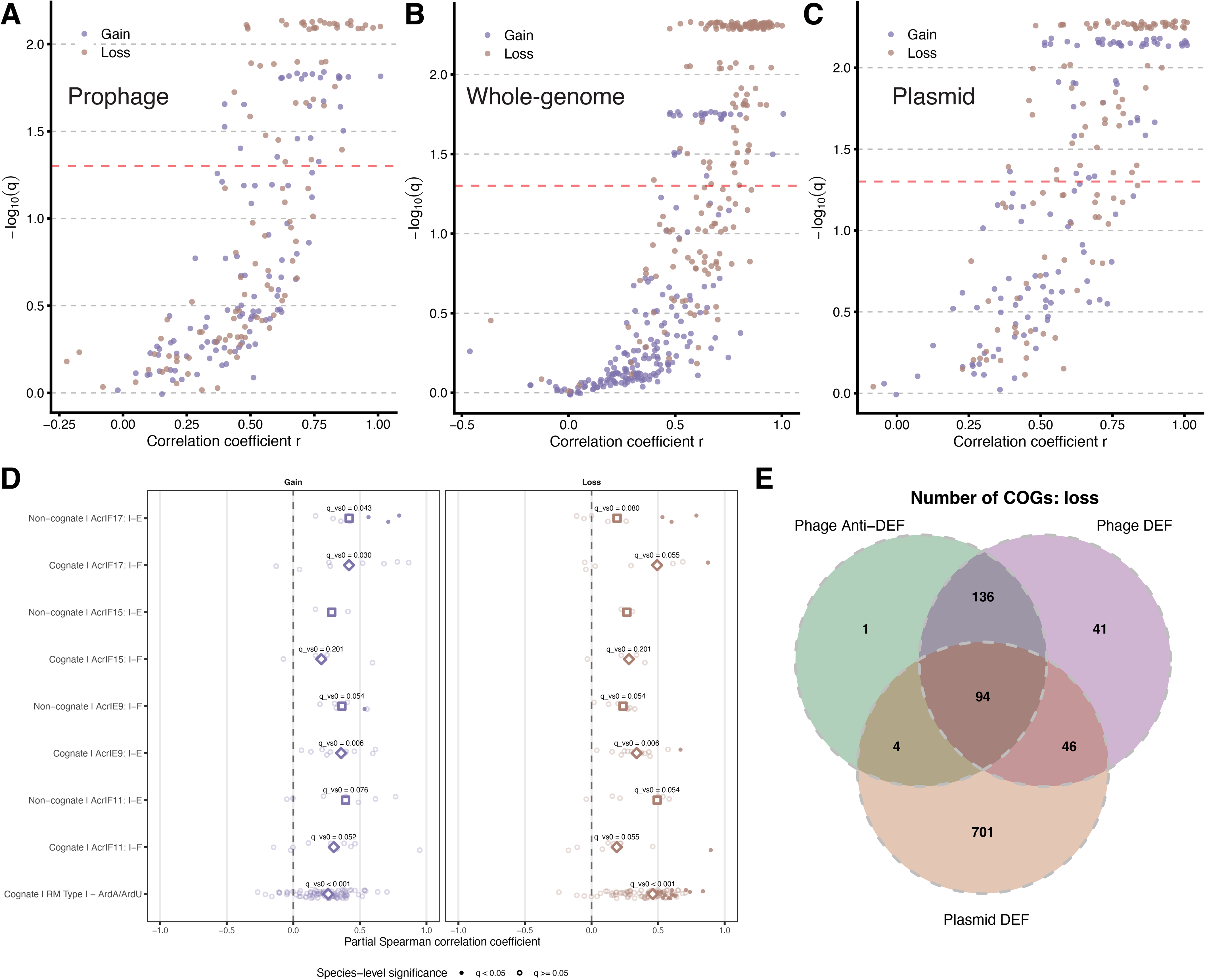
Lineage-level coupled turnover of DEFs and anti-DEFs and overlap with COG-associated gene loss. (A–C) Species-level correlations between the number of gain or loss events per branch involving DEFs and anti-DEFs from prophages (A), whole genomes (chromosome plus MGEs) (B), and plasmids (C). The x-axis represents the partial (branch length-corrected) Spearman correlation coefficient (r), and the y-axis represents statistical significance as −log10(q). Each point represents one species and is colored by event type: gain (purple) or loss (brown). The horizontal dashed line indicates q = 0.05. (D) Species-level partial Spearman correlations for selected matching (that is, antagonistic) DEF–anti-DEF subtype pairs and non-matching pairs belonging to the same broad defense class. Points represent individual species, with symbol fill indicating species-level significance (q < 0.05 or q ≥ 0.05). White diamonds indicate the median correlation coefficient across species. q_vs0 values are the Benjamini–Hochberg false discovery rates for two-sided Wilcoxon signed-rank tests comparing the species-level correlation coefficients against the null hypothesis of zero correlation. (E) Venn diagram showing the overlap of COGs whose loss was significantly correlated with the loss of prophage-associated anti-DEFs, prophage-associated DEFs, or plasmid-associated DEFs. Numbers indicate the number of COGs in each intersection.

Next, we asked whether these associations were specifically observed for matching (that is, antagonistic) DEF and anti-DEF subtype pairs. We examined matching pairs selected from well-represented anti-DEF subtypes with defined DEF targets (Figure 6D). Due to limited sample size, we only conducted these analyses at the whole-genome level. RM Type I–ArdA/ArdU showed positive correlations for both gain (median r = 0.262, n = 92 species, *q* < 0.001) and loss (median r = 0.460, n = 92 species, *q* < 0.001). CRISPR-Cas I-E–AcrIE9 also showed positive correlations for both gain and loss, whereas CRISPR-Cas I-F–AcrIF17 exhibited a significant positive association for gain. It is noteworthy that positive associations were also observed among some non-matching pairs. For example, CRISPR-Cas I-E–AcrIF17 showed a significant positive association for gain (median r = 0.419, n = 8, *q* = 0.043). These association patterns suggest that coupled turnover of DEF and anti-DEF involves not only direct arms races among matching pairs but also broader (non-pair-specific) HGT processes.

Finally, we investigated whether the turnover of MGE-associated DEF and anti-DEF was correlated with the turnover of broader genes. Significant correlations with COG-annotated genes were observed for gain and loss events in both prophage and plasmid regions (Supplementary Figure S6A–C). For gain events, 223 COGs were shared between prophage DEFs and anti-DEFs. Of these, 130 were shared exclusively by the two groups, and 93 were also shared with plasmid DEFs (Supplementary Figure S6D). Similarly, for loss events, 230 COGs were shared between prophage DEFs and anti-DEFs, including 136 that were shared exclusively by these two groups and 94 that were shared with plasmid DEFs (Figure 6E). These overlaps suggest that DEF and anti-DEF turnover may occur as part of broader changes in MGE-associated gene content.

## DISCUSSION

The evolutionary dynamics of bacterial DEFs and anti-DEFs are governed by a complex interplay between the benefits of (anti-)immunity and the costs of maintaining these systems in the genome ^5,28–30^. Our analyses show that genomic background influences the overall flux of DEFs, with the highest stability found in prophages, followed by conjugative and mobilizable MGEs (plasmids, ICEs, and IMEs) and the chromosome (Figure 3A). Despite these differences, the gain-to-loss and expansion-to-reduction ratios do not significantly differ between chromosomal and MGE-associated DEFs (Figure 3B). DEF and anti-DEF gain and loss are evolutionarily coupled across bacterial lineages (Figure 6A-C), although anti-DEFs generally exhibit faster loss and copy-number changes than DEFs (Figure 4-5).

The higher turnover of chromosomal DEFs relative to chromosomal non-defense genes (Figure 2C) likely reflects the substantial fitness cost of defense systems and their short-term benefit in the context of host-parasite arms races ^31,32^. Mechanistically, turnover of chromosomal DEFs could result from processes that expose defense *loci* to horizontal gene transfer even when they are not part of MGEs. Chromosomal defense systems can be mobilized by lateral transduction, whereby prophages or phage-inducible chromosomal islands transfer adjacent chromosomal regions, including defense loci ^33,34^. In addition, some DEFs classified here as chromosomal may reside within integrated MGEs that are not recognized by current annotation methods. More broadly, defense systems are known to cluster in defense islands and other evolutionarily dynamic regions of bacterial chromosomes enriched in recently acquired genes and mobile elements ^4,35^. Such genomic organization may place apparently chromosomal DEFs within regions of elevated genomic plasticity and horizontal gene flux, providing a possible explanation for their rapid turnover despite their chromosomal location.

Overall, DEF turnover was highest on chromosomes, intermediate on plasmids/ICEs/IMEs, and lowest on prophages (Figure 3A). The relatively low turnover of prophage-associated DEFs was consistent with their lower loss and copy-number dynamics compared to the surrounding prophage gene pool (Figure 2C). The lower loss of prophage-associated DEFs relative to other prophage genes suggests that these loci may be selectively maintained. This pattern is also compatible with prophage domestication, whereby beneficial prophage-encoded functions may be preferentially retained as other phage genes are progressively lost or degraded ^36^. Strong retention of prophage-associated DEFs may also reflect the benefits that these systems provide to the resident MGE, for example in competition with other MGEs ^37,38^, although our analyses cannot distinguish benefits to the bacterial host from those to the MGE.

Our findings raise the intriguing hypothesis that chromosomal defense islands could be even more plastic than genuinely mobile regions of the genome, such as prophages and ICEs/IMEs. Besides the lateral transduction mechanisms discussed above, it is possible that insertion sequences and other transposable elements in the proximity of defense islands contributed to a greater genomic instability in these regions. A complementary explanation for the differential DEF retention patterns observed among MGEs and chromosomal regions is that what we identified as MGE-associated DEFs and chromosomal DEFs actually represent different stages of the MGE acquisition-domestication-erosion cycle. From that perspective, domestication of a prophage (or ICE/IME) would first lead to the loss of core viral genes (or integrases and relaxases in the case of ICEs/IMEs). Once the mobile genetic context had been completely eroded, the remaining DEFs would appear as chromosomal and their eventual loss would contribute to the observed loss rate of chromosomal (not MGE-associated) DEFs.

Recent theoretical work predicted that, in a bacterial population in which DEFs can be transferred between MGEs and the chromosome, the loss of chromosomal DEFs would be positively selected over the loss of the same DEFs in MGEs, in what would represent a case of defense outsourcing from the chromosome to symbiotic MGEs ^38^. This study provides indirect empirical support to the defense outsourcing hypothesis by showing that DEFs are more rapidly lost from the chromosome than from MGEs. Note, however, that we did not verify whether MGE-associated DEFs correspond to complete functional systems: Some may represent compact or reduced derivatives of larger defense systems, or defense-system components co-opted for inhibitory functions, as documented for CRISPR–Cas systems ^39–41^. Direct analyses of system composition and function will be required to determine the functional status and evolutionary origin of these MGE-associated DEFs.

The higher loss and copy-number turnover of anti-DEFs may reflect several non-exclusive evolutionary processes that also shape DEF evolution. These include Red Queen-like arms-race dynamics driven by reciprocal adaptation between immunity and anti-immunity, as well as transient selective benefits that depend on phage–host or MGE–host interactions ^14,23,27,30,42^. Anti-DEFs may be especially sensitive to such changes because their benefit depends on the presence and identity of the defense systems they target; when the relevant target is absent or changes, an anti-DEF may become dispensable and be lost because of the cost of maintaining a non-essential function ^28,30,43^. Differences among anti-DEF families may therefore reflect variation in target specificity, selective benefits, maintenance costs, and ecological interactions. Mobility-driven gene flux may additionally contribute to anti-DEF turnover, although the higher loss and copy-number turnover of anti-DEFs compared with DEFs within the same genomic compartments suggests that differences in genomic distribution alone are insufficient to explain the observed trends. As in the case of DEFs, MGE domestication could play a role in anti-DEF dynamics: Anti-DEFs are conceivably less beneficial to the host than DEFs and, as a result, they are likely to be eroded earlier in the domestication process, resulting in higher effective loss rates.

Coupled turnover of DEFs and anti-DEFs occurs at the whole-genome level and involves prophage-associated and plasmid-associated genes (Figure 6A–C). Analysis of specific subtype pairs further revealed positive correlations for both matching and non-matching DEF/anti-DEF pairs (Figure 6D). For matching pairs, correlated turnover is consistent with ecological coupling, whereby the presence of a defense system in a microbial community drives selection for its inhibitor among the local MGEs ^30,42,44^. Ecological coupling could involve sequential transfer of DEF and anti-DEF from separate MGEs rather than physical co-transfer of the two systems on the same MGE. Nevertheless, at evolutionary timescales, both events may appear simultaneous. Although the maintenance of DEF and anti-DEF in the same genome may seem paradoxical, co-occurrence of CRISPR-Cas and antagonistic anti-CRISPR (Acr) has been widely reported ^45,46^. Because prophage-encoded Acrs are often found in genomes with self-targeting CRISPR spacers, it has been proposed that Acr protects prophages and adjacent regions of the genome from self-targeted degradation ^47^. Whether the presence of the Acr brings any benefit to the host or just to the prophage, and for how long CRISPR-Cas and Acr can be jointly maintained in the genome are still open questions. For non-matching DEF/anti-DEF pairs, the observed correlations could reflect separate acquisitions in a relatively short time window or exposure to MGEs carrying both DEFs and anti-DEFs (of different types) ^14,37,48^. Although we did not directly assess the physical linkage of DEFs and anti-DEFs in the same MGEs, we observed that turnover of prophage-encoded DEFs and anti-DEFs, as well as plasmid-encoded DEFs, is associated with acquisition and loss of a substantial set of shared COG families (Figure 6E and Supplementary Figure S6). This observation supports the hypothesis that concerted dynamics of MGE-associated DEFs and anti-DEFs reflects broader changes in MGE regions ^36,49^. In sum, the observed coupling between DEFs and anti-DEFs seems to emerge not only from ecological interactions between antagonistic systems but also from background turnover of MGEs that encode non-matching DEF/anti-DEF systems.

Although we conducted comprehensive analyses, statistical power may be limited for species represented by relatively few genomes, particularly when few DEF or anti-DEF OGs are available. Increased genome sampling of these underrepresented species would likely improve comparisons among DEF or anti-DEFs from different genomic regions. The cognate and non-cognate analyses were also limited to a small number of well-represented DEF–anti-DEF pairs because many anti-DEF subtypes were rare. With more data, additional pairs could be tested, and the relative strength of cognate and non-cognate correlations may differ from that observed here. Furthermore, our analysis was restricted to the *Enterobacteriaceae* family. Although this family is highly relevant for understanding HGT, defense turnover dynamics may differ in other bacterial lineages with distinct MGE ecologies.

## METHODS

### Genome Collection and Quality Filtering

We parsed the Genome Taxonomy Database (GTDB; https://gtdb.ecogenomic.org) release 226 ^50^ to identify genomes belonging to Enterobacteriaceae. We applied stringent genome-quality filters, requiring completeness >99%, contamination <1%, mean contig length >5,000 bp, and fewer than 500 contigs. To reduce computational cost, species with more than 50 genomes were subsampled to retain at most 50 genomes. Only species with at least 10 genomes were considered for further analysis. The genomes that passed these filters were downloaded from the NCBI FTP site (https://ftp.ncbi.nlm.nih.gov).

After applying these criteria, 7,059 complete or nearly complete genomes belonging to 220 Enterobacteriaceae species (sensu GTDB) were included in the analysis (Supplementary Table S1).

### Gene Prediction and Functional Annotation

Bacterial genes were predicted using pyrodigal-gv v0.3.2 ^51^ integrated in geNomad. Prophages and plasmids were identified using geNomad v1.11.0 with database v1.9 and default parameters ^52^. Because *Enterobacteriaceae* contain chromosome-plasmid hybrids (chromids) that can complicate plasmid assignment, Platon v1.7 ^53^ was used to further cross-validate plasmid predictions. Genes were classified as plasmid-associated only if they were identified by both tools. Phage satellites were identified using SatelliteFinder v0.9.1 ^54^. ICEs and IMEs were identified using ICEfinder-opt v1.0.0 (https://github.com/guogenglin/icefinder-opt, accessed 16 April 2026), an optimized local version of ICEfinder2 ^55^ that supports both complete and draft genome assemblies. To reduce potential false-positive ICE/IME assignments arising from plasmid-derived sequences, ICE/IME predictions were cross-checked against the plasmid contigs identified above. Predicted ICEs/IMEs located outside these plasmid contigs were retained, whereas those located on plasmid contigs were retained only when supported by four integration-associated features, including *attL*, *attR*, *att* annotations, and nearby tRNA genes. As a validation of this procedure, ICEfinder-opt predictions were compared with those obtained using the original ICEfinder2 implementation in complete genomes. Among 1,228 successfully processed genomes, 992 of 1,100 ICEfinder-opt predictions and 1,014 of 1,065 original ICEfinder2 predictions overlapped at the genomic-coordinate level.

DEFs and anti-DEFs were identified using DefenseFinder version 2.0.1 ^7,23^. Pan-genomes were constructed independently for each species using the orthology-based clustering method implemented in OrthoFinder v2.5.5 ^56^, which was selected based on previously published guidelines ^57^.

For COG functional annotation, we constructed HMM profiles from the COG 2024 database accessed on 20 February 2025 ^58^. Protein sequences belonging to each COG family were aligned using MAFFT v7.526 ^59^ with the *--auto* option and trimmed using ClipKIT v2.6.1 ^60^ in smart-gap mode. HMM profiles were generated for each COG family using *hmmbuild* from HMMER v3.4 (http://hmmer.org) with default parameters. Predicted proteins from each genome were searched against the resulting COG HMM database using *hmmsearch* with an E-value cutoff of 1 x 10^-5^, followed by a more stringent filter requiring a bitscore > 50.

### Gene Family Evolution Modeling Using Birth–Death Processes

Orthologous gene families (OGs) and phylogenomic trees for each species were generated with OrthoFinder v2.5.5. Gene gain and loss dynamics were quantified using the phylogenetic birth-death model implemented in Count v10.04 ^61^, providing gene family copy-number profiles and species trees as inputs. The software Count uses a probabilistic framework to model the evolution of gene family sizes along phylogenetic trees as a continuous-time Markov process. The model accounts for three fundamental processes: gene gain (κ), gene loss (μ), and gene duplication (λ). A gene family of size *n* can reduce its copy number at a rate *n*μ due to individual gene losses and increase through duplications at a rate *n*λ. Moreover, a gene family that is initially absent can be gained at a rate κ through horizontal transfer or other acquisition mechanisms. For each species dataset, Count performs maximum likelihood optimization to estimate model parameters from discretized gamma distributions, with the number of discrete gamma categories given by the user. As recommended by the Count user’s guide and for consistency with previous publications ^9,62^, we implemented an iterative search strategy starting with 1 discrete gamma category for all the parameters and iteratively increasing the number of discrete gamma categories up to 4. Specifically, we changed the number of discrete gamma categories in a stepwise cyclic manner, increasing it in one unit for the loss rate, then the duplication rate, then the gain rate, and back to the loss rate, until reaching 4 discrete gamma categories for all the parameters in 10 rounds of optimization. Each round used the previous round’s results as starting values. This stepwise design and the upper bound of 4 discrete gamma categories help avoid overfitting and maintain numerical stability during likelihood optimization ^61^. Optimization stability across successive rounds was assessed using likelihood trajectories across the 10 optimization rounds (Supplementary Figure S2). Count options for maximum likelihood estimation were set to 200 maximum paralogs, optimization tolerance of 0.1, and 2 minimum lineages.

After parameter optimization, we calculated posterior probabilities for ancestral gene family states and evolutionary events using the Posteriors module of Count. This reconstructs the evolutionary history of each gene family, providing probabilistic estimates for gain, loss, expansion, and reduction events along each branch. Count analyses were conducted independently for each species dataset. Of the 220 species initially selected, 213 species comprising 6,907 genomes yielded usable Count results and were included in downstream evolutionary rate analyses.

### Calculation of Branch-Level and OG-Level Evolutionary Rates

Orthologous gene families were classified based on the genomic location of their member genes. An OG was assigned to an MGE class when ≥90% of its member genes were located within prophages, plasmids, ICEs/IMEs, or phage satellites, with the threshold applied independently to each MGE class. OGs were classified as chromosomal when <10% of their member genes were located within any annotated MGE region. OGs that satisfied neither criterion were considered ambiguous and were excluded from comparisons.

Plasmid- and ICE/IME-associated OGs were subsequently pooled into a single plasmid plus ICE/IME (*pl+i*) category because both classes of MGEs share the same transfer mechanisms and their DEF turnover rates were generally similar. Prior to merging, within each species and evolutionary event type, plasmid-associated and ICE/IME-associated DEF OGs were compared using two-sided Mann–Whitney U tests, followed by Benjamini–Hochberg correction. Significant differences were detected in only 2 of 22 species for gain and 1 of 22 for reduction, and in none for loss or expansion.

For the descriptive UpSet plots in Figures 2A–B, additional combined categories were used to display OGs whose member genes were distributed across multiple genomic compartment classes and collectively reached the 90% threshold. These combined categories were used only for visualization and were excluded from downstream analyses.

For genomic compartment turnover comparisons, gene families were classified as DEF if they contained at least three defense-annotated genes or if ≥20% of their member genes were annotated as defense genes. Similarly, gene families were classified as anti-DEF if they contained at least three anti-defense-annotated genes or if ≥20% of their members carried anti-defense annotations.

Branch lengths were extracted from the species trees used as input for Count and matched to the corresponding posterior event estimates. Posterior event estimates reported as NaN were excluded from downstream rate calculations. For each retained OG-event combination, posterior expectations of gain, loss, expansion, and reduction events were summed separately across all branches and divided by the total tree length of the corresponding species. Thus, branches with zero posterior events contributed zero to the numerator but were retained in the denominator. Zero-length branches were excluded from rate calculations. Evolutionary rates were calculated for each event type and OG as:

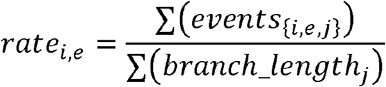

where *events_{i,e,f}_* is the posterior expected number of events of type *e* for OG *i* on branch *j*, and *branch_length_{j}_* is the corresponding branch length in substitutions per site.

### Comparison of DEF and Non-Defense Gene Turnover

To determine whether DEF turnover rates differed from the background dynamics of the genomic regions in which they were located, DEF OGs were compared with non-defense (ND) OGs from the same genomic compartment. Comparisons were performed independently for each species and evolutionary event type using OG-level gain, loss, expansion, and reduction rates. Two-sided Mann–Whitney U tests were performed only when both groups contained at least three OGs. *P*-values were adjusted using the Benjamini–Hochberg procedure separately within each evolutionary event and comparison type.

Effect sizes were quantified using the rank-biserial correlation derived from the Mann–Whitney U statistic. A threshold of (|r| ≥ 0.147) was used as the primary small-effect-size criterion ^63,64^. To assess the robustness of the conclusions to the choice of effect-size threshold, the analyses were repeated using more stringent thresholds of (|r| ≥ 0.33) and (|r| ≥ 0.474), corresponding to medium and large effects ^63,64^, respectively. To assess the overall direction of effects across species, two-sided exact binomial tests were used to compare the numbers of species with positive and negative effect sizes exceeding each threshold, excluding species with smaller effects. Overall differences between DEF and ND turnover were additionally evaluated using GLMMs as described below.

### Generalized Linear Mixed-Effects Model Analyses of Gene Flux

Generalized linear mixed-effects models (GLMMs) were fitted separately for gain, loss, expansion, and reduction using glmmTMB ^65^ v1.1.14 with a Gamma distribution and log link. The summed posterior expectation of each evolutionary event for each species–OG combination was used as the response, with the logarithm of the total species-tree branch length included as an offset. (Note that, because of the log link function, introducing the logarithm of the total species-tree branch length as an offset guarantees that the total number of events will be proportional to the total evolutionary time, as logically expected.) Models were fitted only when each fixed-effect level was represented in at least three species.

To compare DEFs with ND genes within the same genomic compartment, separate models were fitted for chromosome, prophage, and plasmid plus ICE/IME backgrounds:

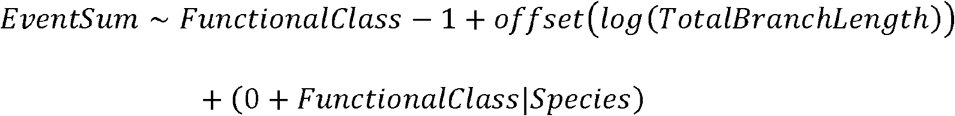

where *FunctionalClass* represented DEF or ND.

To compare DEF turnover among genomic locations, the model was:

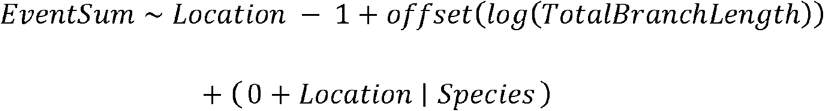

where *Location* included chromosome, prophage, and plasmid plus ICE/IME; phage satellite-associated DEFs did not meet the minimum representation criterion above.

To compare DEFs and anti-DEFs within the same genomic compartment, separate models were fitted for chromosome, prophage, and plasmid plus ICE/IME backgrounds:

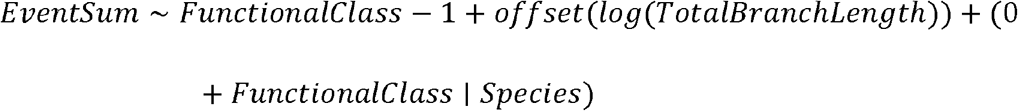

where *FunctionalClass* represented DEF or anti-DEF. OGs annotated as both DEF and anti-DEF were excluded.

For models in which the random-slope structure resulted in overparameterized or otherwise unsuitable fits, a random-intercept structure, (1│*species*), was used instead. This applied to the gain comparisons among DEF genomic locations, between cDEF and cND genes, between pDEF and pND genes, and between pDEFs and pAnti-DEFs. A model without’ random effects were also tested, and the random intercept model was selected based on AIC.

Pairwise contrasts were estimated using *emmeans* v2.0 ^66^ and reported as rate ratios with 95% confidence intervals. *P*-values were adjusted using the Benjamini–Hochberg procedure separately within each analysis and evolutionary event type.

### GLMM Analysis of Gain/Loss and Expansion/Reduction Ratios

To compare the relative balance of DEF turnover between chromosomal and MGE backgrounds, gain/loss and expansion/reduction ratios were analyzed using GLMMs. To ensure within-species comparison of genomic backgrounds, analyses were restricted to species containing DEF OGs in both chromosomal and MGE categories. Separate models were fitted for gain versus loss and expansion versus reduction using a Tweedie distribution with a log link:

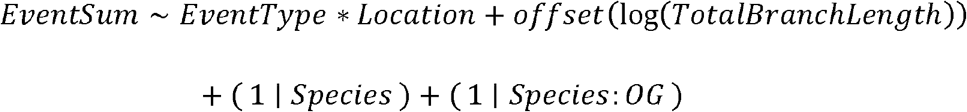

where *EventType* was a categorical variable with levels gain and loss in the gain/loss model, and expansion and reduction in the expansion/reduction model; *Location* was a categorical variable indicating chromosome or MGE. The *EventType × Location* interaction was exponentiated to obtain the MGE-to-chromosome ratio of rate ratios. *P*-values for the two ratio-of-ratios tests were adjusted using the Benjamini–Hochberg procedure.

### Comparative Analysis of DEF and Anti-DEF Turnover

Rate differences between DEFs and their cognate anti-DEFs were quantified by pairing each DEF class with its corresponding anti-DEF class within each species and comparing their median rates. OGs annotated as both DEF and anti-DEF were excluded as described above. Evolutionary rates (gain, loss, expansion, and reduction) were retrieved per OG per species, and system-level turnover was summarized as the median rate across all OGs within each system. For every DEF versus anti-DEF pair and event type, paired differences in median turnover and median fold changes between DEF and anti-DEF were calculated. Paired Wilcoxon signed-rank tests were performed when at least three species observations were available for a given system pair, followed by Benjamini–Hochberg correction within each event type.

### Correlated Evolutionary Event Analysis Using Partial Spearman Correlations and Phylogenetic Brownian Motion

Correlated evolutionary events were assessed by testing whether DEFs, anti-DEFs, and COG-annotated gene families exhibited similar gain or loss patterns across phylogenetic branches within each species. For DEF versus anti-DEF comparisons, branch-specific gain and loss events were summed separately across DEF and anti-DEF gene families for whole-genome, prophage- and plasmid-specific analyses. For comparisons with COG-annotated gene families, correlations were tested within the same genomic element class: prophage-associated DEFs and anti-DEFs were compared with prophage-associated COG families, and plasmid-associated DEFs were compared with plasmid-associated COG families. A gene family was assigned a COG ID if ≥75% of that family shared the same COG annotations. Branch-specific gain and loss events were obtained from Count posterior estimates. For these analyses, DEF and anti-DEF OGs were identified directly from DefenseFinder annotations without applying the previously described thresholds.

Associations between branch-specific events were quantified using partial Spearman correlations controlling for branch length. The correlation coefficient was calculated as the Pearson correlation between the residuals obtained after regressing each rank-transformed event measure on rank-transformed branch length. Both terminal and internal branches were included, and comparisons involving fewer than eight branches were excluded. Phylogenetic Brownian motion (BM) simulations were used to generate *null* distributions for significance testing, with 500 simulations per comparison. Two-sided empirical *P*-values were calculated from the simulated correlation distributions and adjusted using the Benjamini–Hochberg procedure within each event type. Species trees were midpoint-rooted before analysis, and Brownian-motion traits were simulated using fastBM, and ancestral states were reconstructed using fastAnc, both implemented in the phytools package v2.5.2 ^67^ for R v4.5.1.

We then assessed cognate DEF and anti-DEF coupled turnover at the subtype level. The cognate DEF and anti-DEF pairs were selected based on abundance and distribution, and non-cognate pairs were predefined within the same broad defense class. For each species and subtype pair, partial Spearman correlations were calculated as described above. This yielded one correlation coefficient for each species, event type, genomic compartment, and subtype pair. For each subtype pair, the median species-level correlation coefficient was used to summarize the direction and magnitude of the association, while a two-sided Wilcoxon signed-rank test was used to assess whether the distribution of correlation coefficients across species differed from zero. *P*-values were adjusted using the Benjamini–Hochberg procedure separately for cognate and non-cognate pairs within each genomic compartment and event type.

Significantly correlated COG-annotated gene families were categorized according to COG functional categories. Individual COG categories were mapped to four major process groups: information storage and processing, cellular processes and signaling, metabolism, and poorly characterized functions.

## ACKNOWLEDGEMENTS

This work was funded by the China Postdoctoral Science Foundation (grant no. 2025M783825 to Yang Liu) and by MICIU/AEI/10.13039/501100011033 and ERDF/EU (PGE) ‘A way of making Europe’ (grants PID2019-106618GA-I00 and CNS2023-145430 to Jaime Iranzo). J.B. acknowledges financial support from national funds through FCT – Fundação para a Ciência e a Tecnologia under contract no. 2023.15056.TENURE.054. J.B. also acknowledges financial support from national funds through the FCT - Foundation for Science and Technology projects UID/50016/2025 (https://doi.org/10.54499/UID/50016/2025) and LA/P/0076/2020 (https://doi.org/10.54499/LA/P/0076/2020). We would also like to thank two anonymous reviewers for their comments on a previous version of this paper.

## DECLARATION OF GENERATIVE AI

ChatGPT and Gemini were used to assist with code drafting and language editing. All generated code was reviewed by the authors, and all manuscript text was edited by the authors, who take full responsibility for the final content.

## DECLARATION OF COMPETING INTERESTS

The authors declare no competing interests.

## ETHICS APPROVAL

There are no ethical concerns in this study.

## DATA AVAILABILITY

The data and analysis scripts for the evolutionary-rate comparisons and phylogenetic correlation analyses are available on Figshare at https://doi.org/10.6084/m9.figshare.30737930. Genome accession numbers are provided in Supplementary Table S1, and figure source data are provided with the article.

## AUTHOR CONTRIBUTIONS

**Y.L.** conceived the study, performed bioinformatic analyses, acquired funding, and drafted and revised the manuscript. **S.D.A.** performed bioinformatic analyses for the original submission and contributed to drafting and revising that version of the manuscript. **J.B.** contributed to the identification and interpretation of ICEs and IMEs and revised the manuscript. **J.I.** contributed mathematical and statistical methodology, refined the analytical framework and interpretation, and substantially revised the manuscript. All authors approved the final manuscript.

## Notes

### Competing Interest Statement

The authors have declared no competing interest.

### Summary of Updates

The revision included the identification of additional classes of mobile genetic elements (phage satellites and integrative and conjugative/mobilizable elements), the recalculation of evolutionary rates, and the implementation of more sophisticated statistical methods (generalized linear mixed-effects models) to verify the robustness of the results. As a result, we identified statistically significant trends that were not detected in the original analysis, which required a thorough rewrite of the Results and Discussion sections.

