## Supplemental Figure for "Co-evolutionary dynamics of defense systems and anti-defense genes in chromosomes and mobile genetic elements of *Enterobacteriaceae*"

### Supplementary Figure Legends

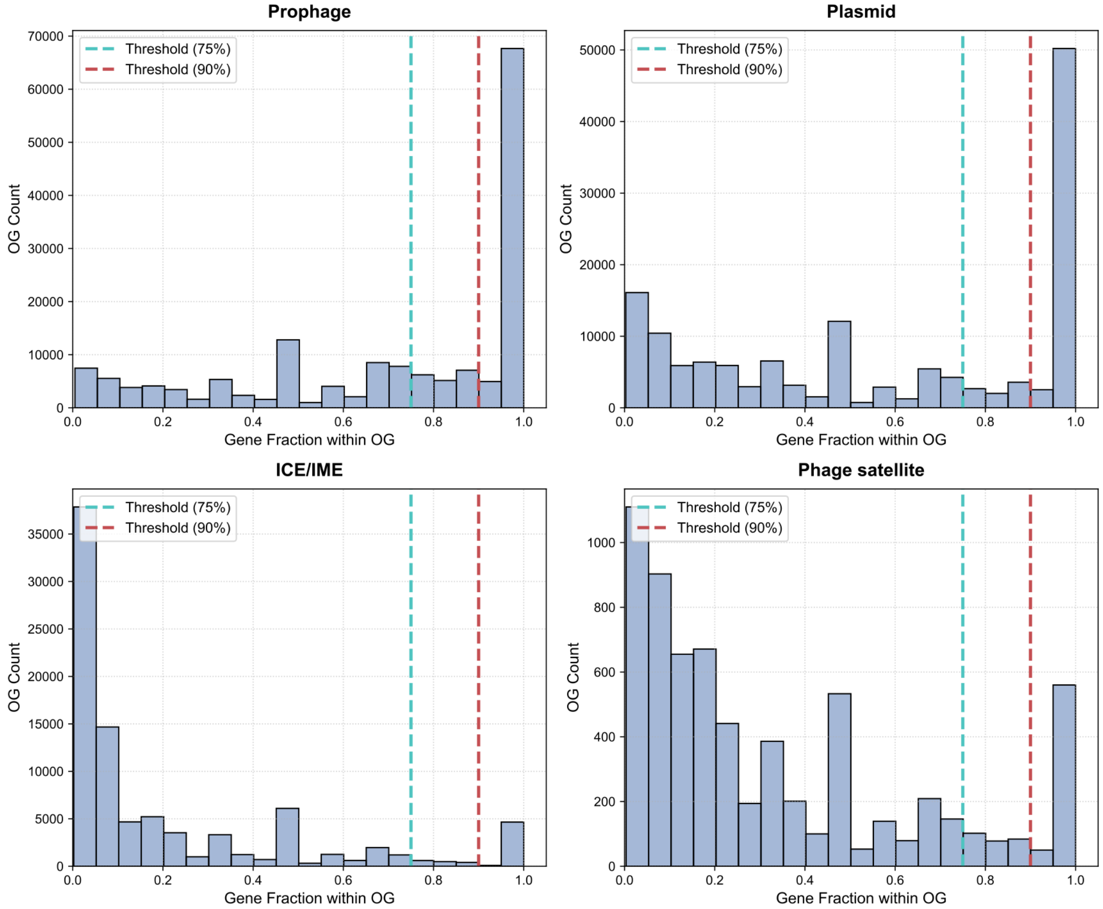


**Figure S1. Distribution of mobile genetic element (MGE)-associated gene fractions within orthologous groups (OGs).** (A) prophages, (B) plasmids, (C) ICEs/IMEs, and (D) phage satellites. The x-axis represents the proportion of genes within an OG located in the corresponding MGE category, and the y-axis represents the number of OGs. Dashed lines indicate the 75% and 90% thresholds used to evaluate MGE-associated OG classification.

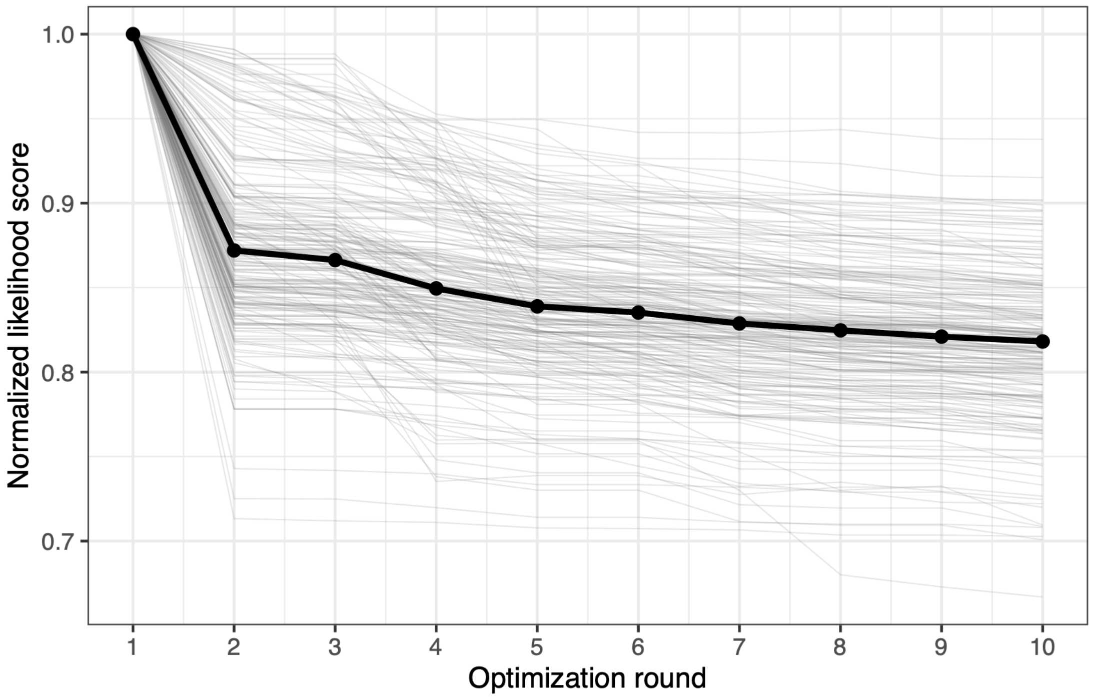


**Figure S2. Likelihood trajectories across stepwise optimization rounds in Count.** For each species, the likelihood score at each optimization round was normalized to the likelihood score obtained in the first round. Thin grey lines represent individual species and the black line represents the median across all species.

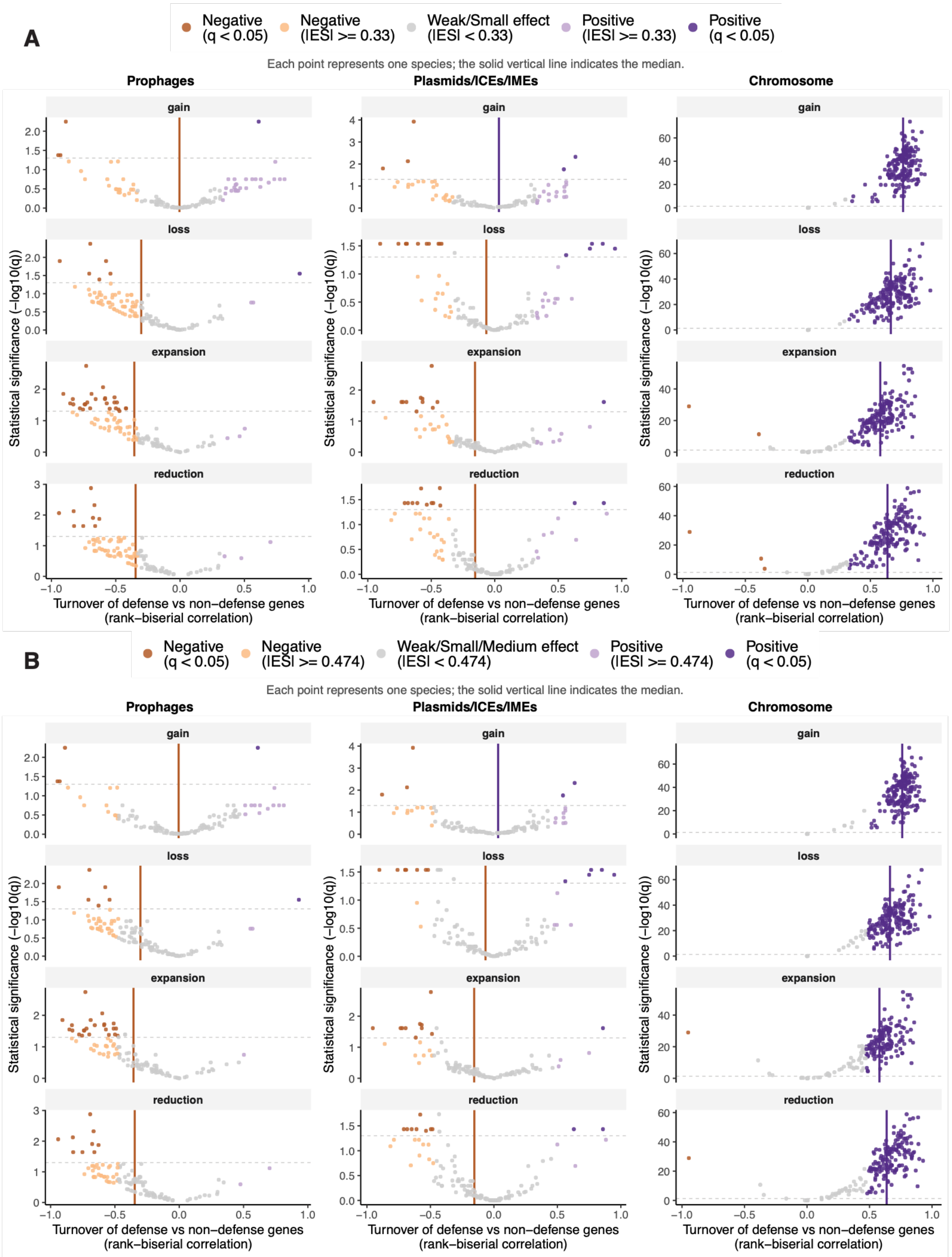


**Figure S3. Differences in evolutionary rates between DEFs and non-defense genes (ND) located within the same genomic element class using more stringent effect-size thresholds.** (A) Effect-size threshold of 0.33. (B) Effect-size threshold of 0.474. Comparisons are shown for prophage genes (pDEF vs pND), plasmid plus ICE/IME genes (pl+iDEF vs pl+iND), and chromosomal genes (cDEF vs cND). Each point represents one Enterobacteriaceae species. Colors indicate the direction and statistical/effect-size support of the difference. Solid vertical lines indicate the median effect size across species. ES, effect size.

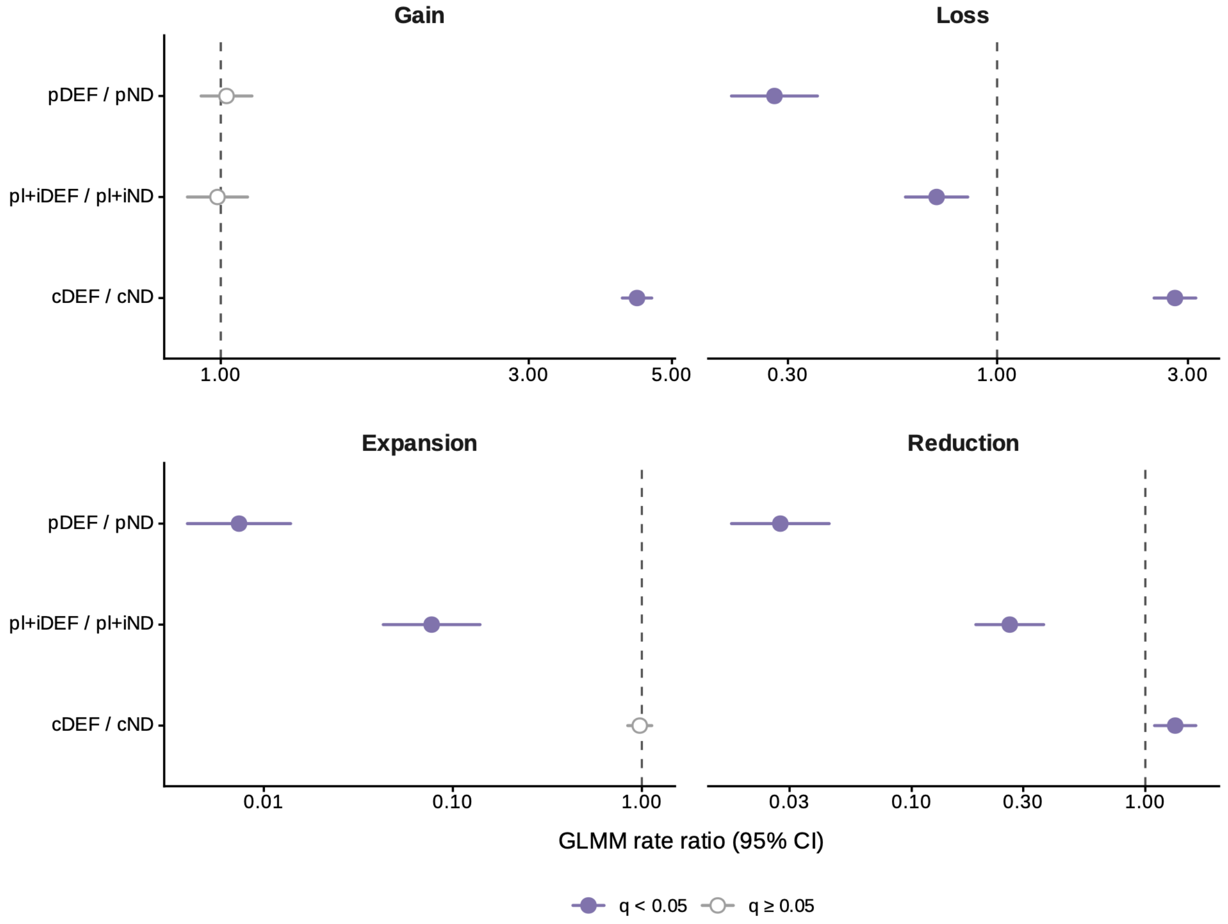


**Figure S4. GLMM-based comparison of evolutionary turnover between DEFs and non-defense genes across genomic compartments.** Comparisons of DEF and non-defense gene evolutionary rates within prophages (pDEF/pND), plasmids plus ICEs/IMEs (pI+iDEF/pI+iND), and chromosomes (cDEF/cND) for gain, loss, expansion, and reduction events. Points represent estimated rate ratios from generalized linear mixed models (GLMMs), and horizontal lines indicate 95% confidence intervals (CIs). The dashed vertical line indicates a rate ratio of 1. Filled points indicate $q < 0.05$ and open points indicate $q \geq 0.05$.

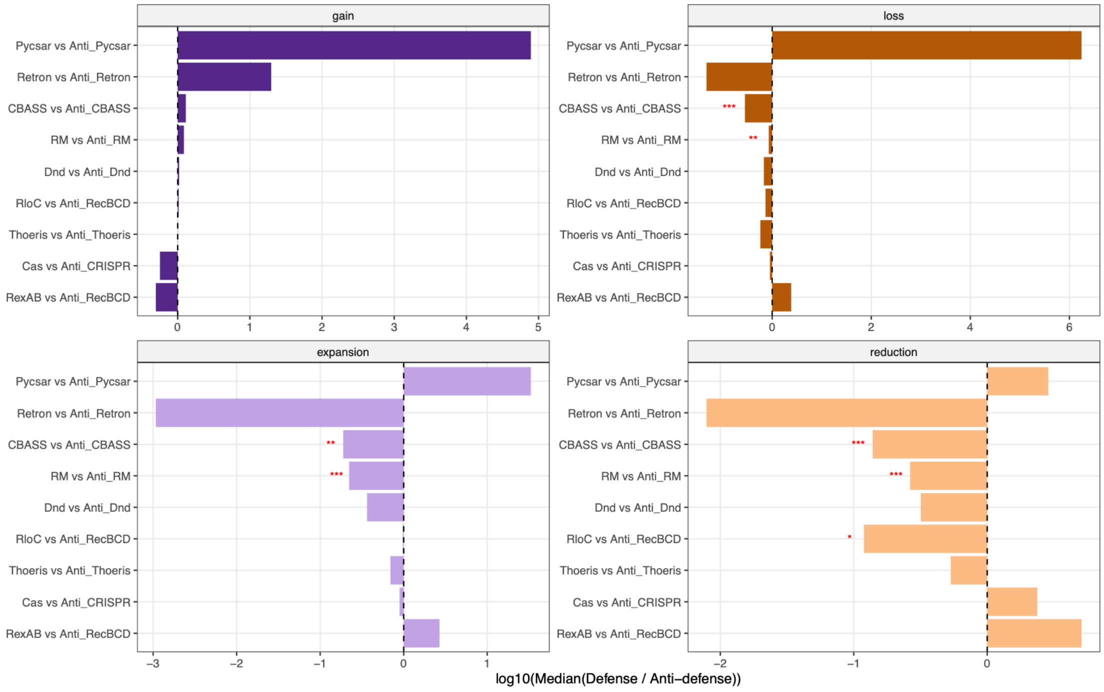


**Figure S5. Relative differences in evolutionary rates between specific DEFs and anti-DEFs.** Bar charts show the magnitude of rate differences for specific DEF and anti-DEF pairs. The x-axis represents the log10-transformed median of species-level DEF-to-anti-DEF rate ratios. Positive values indicate higher rates in the DEF, whereas negative values indicate higher rates in the anti-DEF. Red asterisks indicate statistically significant differences based on paired Wilcoxon signed-rank tests after multiple-testing correction (* $q < 0.05$, ** $q < 0.01$, *** $q < 0.001$).

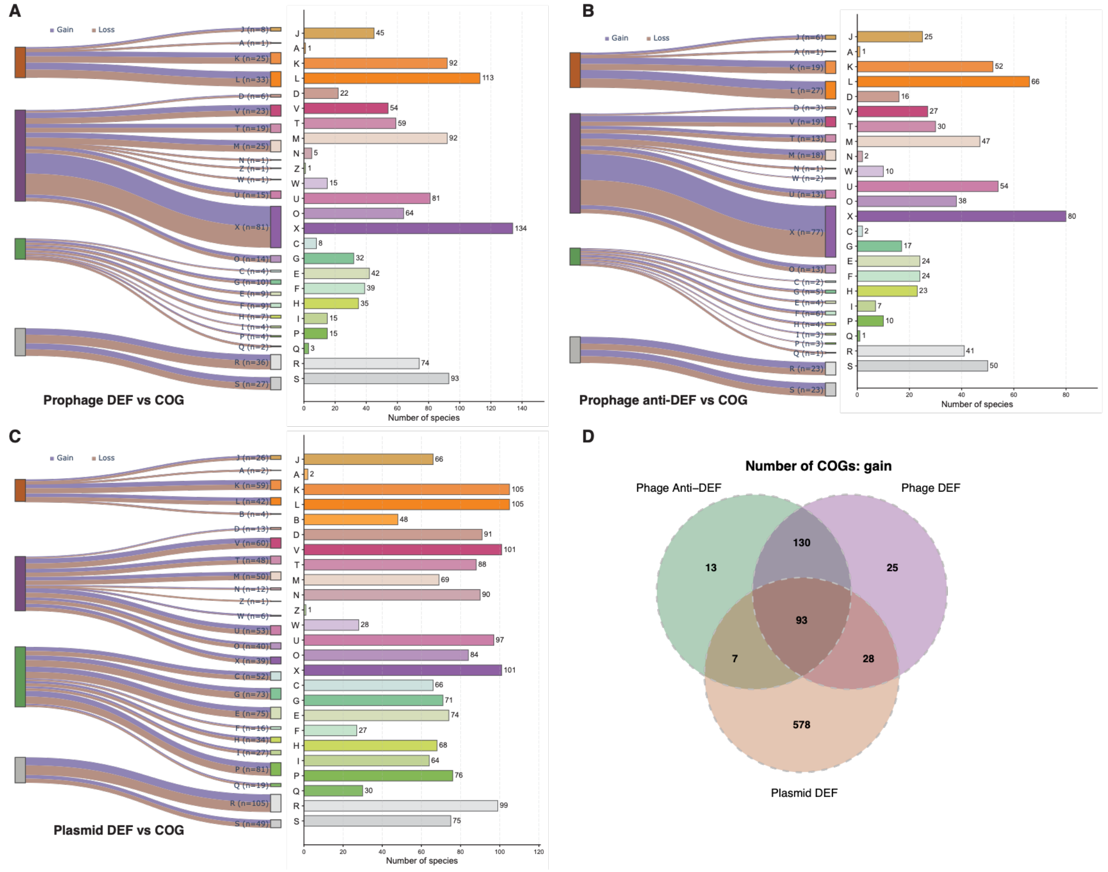


**Figure S6. Functional gene categories showing coupled turnover with prophage-associated DEFs and anti-DEFs and plasmid-associated DEFs.** (A–C) Significant correlations between COG-annotated gene families and prophage-associated DEFs (A), prophage-associated anti-DEFs (B), or plasmid-associated DEFs (C). Sankey diagrams show the distribution of significantly correlated COGs across functional categories, and adjacent bar plots show the number of species exhibiting significant correlations for each COG category. Colors of the flows indicate gain and loss events, whereas flow width and the numbers in parentheses following each COG category label indicate the number of COGs identified within that category. The four major process groups shown on the left, from top to bottom, are information storage and processing, cellular processes and signaling, metabolism, and poorly characterized functions. (D) Venn diagram showing the overlap of COGs whose gain was significantly correlated with the gain of prophage-associated anti-DEFs, prophage-associated DEFs, or plasmid-associated DEFs. Numbers indicate the number of COGs in each intersection. COG category abbreviations are as follows: A: RNA processing and modification; B: Chromatin structure and dynamics; C: energy production and conversion; D: cell cycle control, cell division, chromosome partitioning; E: amino acid transport and metabolism; F:

nucleotide transport and metabolism; G: carbohydrate transport and metabolism; H: coenzyme transport and metabolism; I: lipid transport and metabolism; J: translation, ribosomal structure and biogenesis; K: transcription; L: replication, recombination and repair; M: cell wall/membrane/envelope biogenesis; N: cell motility; O: posttranslational modification, protein turnover, chaperones; P: inorganic ion transport and metabolism; Q: secondary metabolites biosynthesis, transport and catabolism; R: general function prediction only; S: function unknown; T: signal transduction mechanisms; U: intracellular trafficking, secretion, and vesicular transport; V: defense mechanisms; W: extracellular structures; X: mobilome: prophages, transposons; Y: nuclear structure; Z: cytoskeleton.
